# Mitochondrial clearance of Ca^2+^ controls insulin secretion

**DOI:** 10.1101/830323

**Authors:** N Vishnu, A Hamilton, A Bagge, A Wernersson, E Cowan, H Barnard, Y Sancak, K.J. Kamer, P Spégel, M Fex, A Tengholm, V.K. Mootha, DG Nicholls, H Mulder

## Abstract

Transport of Ca^2+^ from the cytosol to the mitochondrial matrix of insulin-secreting pancreatic β-cells facilitates nutrient-mediated insulin secretion. However, the underlying mechanism is unclear. The establishment of the molecular identity of the mitochondrial Ca^2+^ uniporter (MCU) and associated proteins has allowed mitochondrial Ca^2+^ transport to be modified in intact cells. We examined the consequences of deficiency of the accessory protein, MICU2, in rat and human insulin-secreting cell lines as well as in mouse islets. Glucose-induced mitochondrial Ca^2+^ elevation and inner membrane hyperpolarization were reduced, together with cytosolic ATP/ADP-ratios and insulin secretion. Insulin secretion in *Micu2* knock out mice was attenuated *in vitro* as well as *in vivo*. While KCl-evoked sub-plasmalemmal Ca^2+^ increases were more pronounced, the global cytosolic Ca^2+^ response was, surprisingly, diminished in *MICU2*-deficient cells. These findings were supported by selective inhibition of mitochondrial Ca^2+^ uptake by mitochondrial depolarization. It is concluded that mitochondrial Ca^2+^ transport plays an additional and hitherto unrecognized role in stimulated β-cells by regulating net Ca^2+^ entry across the plasma membrane. This is likely accounted for by clearing of sub-plasmalemmal Ca^2+^ levels by mitochondria located near the plasma membrane.

## INTRODUCTION

Diabetes is a multifactorial metabolic disease that manifests itself as chronic hyperglycemia. There is currently a soaring increase in the prevalence of Type 2 Diabetes (T2D), caused by the global obesity epidemic. T2D involves deficiencies in insulin secretion as well as in effects of the hormone in its target tissues (Tuomi, Santoro et al., 2014). The fundamental stimulus for insulin secretion from pancreatic β-cells is glucose (Seino, Shibasaki et al., 2011). This involves an increase in glucose uptake and subsequent metabolism of the hexose by the β-cells, raising the cellular ATP/ADP ratio, which leads to closure of ATP-dependent potassium (K_ATP_)-channels. Consequently, the plasma membrane is depolarized and voltage-dependent Ca^2+^-channels open, allowing Ca^2+^ influx. The elevated free cytosolic Ca^2+^ concentration ([Ca^2+^]_c_) subsequently provokes exocytosis of insulin granules (Ashcroft, 2005).

The mitochondrial bioenergetic response to altered glucose availability is central to this mechanism (Ashcroft, 2005). Mitochondria have the capacity to take up and buffer intracellular Ca^2+^. The free matrix Ca^2+^ concentration, [Ca^2+^]_mito_, increases when [Ca^2+^]_c_ is elevated, thereby contributing to buffering of the ion in the cell. [Ca^2+^]_mito_ may also affect the tone and frequency of [Ca^2+^]_c_ oscillations (Wiederkehr & Wollheim, 2008), a regulation which may play a crucial role in controlling the pulsatility of insulin release (Gilon, Shepherd et al., 1993). Moreover, [Ca^2+^]_mito_ is required for activation of metabolic enzymes involved in the tricarboxylic acid (TCA) cycle, i.e., the dehydrogenases, as well as in oxidative phosphorylation, i.e., ATP synthase, reactions that are essential for effective ATP generation in pancreatic β-cells, as well as in any other cell (Denton & McCormack, 1986, Wiederkehr & Wollheim, 2008). These circumstances support the proposition that mitochondrial Ca^2+^ import is a critical event in cellular bioenergetics and, in extension, metabolic coupling of insulin secretion. Despite this, precisely how [Ca^2+^]_mito_ contributes to the regulation of insulin secretion is not fully understood.

Mitochondrial Ca^2+^ uptake in most cells occurs via a macromolecular protein complex, known as the calcium uniporter holocomplex (Jhun, Mishra et al., 2016, Sancak, Markhard et al., 2013). MCU forms the central pore-forming subunit of the complex, while MCU regulatory subunit b (MCUb), essential MCU regulator (EMRE), mitochondrial uptake 1 and 2 (MICU1 and MICU2) form the accessory components (Kamer & Mootha, 2015). As part of the Ca^2+^ uniporter holocomplex, MICU1 and MICU2 heterodimerize and act to maintain a threshold for mitochondrial Ca^2+^ uptake, upholding the mitochondrial Ca^2+^ buffering capacity (Ahuja & Muallem, 2014, Kamer, Grabarek et al., 2017, Kamer & Mootha, 2014). It has been demonstrated in INS-1 832/13 insulinoma cells that MICU1 plays a role in metabolic coupling; its knock down leads to reduced mitochondrial Ca^2+^ uptake and inhibition of insulin secretion (Alam, Groschner et al., 2012). However, the function of MICU2 in stimulus-secretion coupling in pancreatic β-cells has not been examined.

To this end, we examined the impact of MICU2-deficiency in insulin-secreting cells and mouse islets. We observed that [Ca^2+^]_mito_ elevation upon high glucose stimulation was significantly reduced by *MICU2*-deficiency, as was the cytosolic ATP/ADP-ratio and glucose-stimulated insulin secretion (GSIS). *Micu2* knock out mice were normoglycemic but released less insulin both *in vivo* and *in vitro*. Counterintuitively, *MICU2*-deficiency impaired high potassium and glucose-induced [Ca^2+^]_c_ elevations. This effect was associated with a transient elevation of sub-plasmalemmal Ca^2+^ levels ([Ca^2+^]_mem_). Our data suggest that mitochondrial Ca^2+^ transport enhances net cellular Ca^2+^ entry, directly or indirectly, and point to a hitherto unknown role of the mitochondrial Ca^2+^ uptake in the control of insulin secretion.

## RESULTS

### Functional Consequences of *MICU2* Knock down in Insulin-Secreting cells

To understand the role of MICU2 in stimulus-secretion coupling in pancreatic β-cells, we silenced *MICU2* in INS-1 832/13 cells, a rodent-derived insulinoma cell line (Hohmeier, Mulder et al., 2000), and in EndoC-βH1 cells, a human embryonic insulin-secreting cell line (Andersson, Valtat et al., 2015). Our experiments revealed that after 48 or 72 hours, small interfering (si) RNA treatment was sufficient to significantly reduce expression of *MICU2* at the mRNA and protein level in both cell lines (Figure 1A-D). Notably, *MCU* and *MICU1* expression were unaffected by *MICU2* silencing (Supplementary Figure 1 A-B).

**Figure 1.**
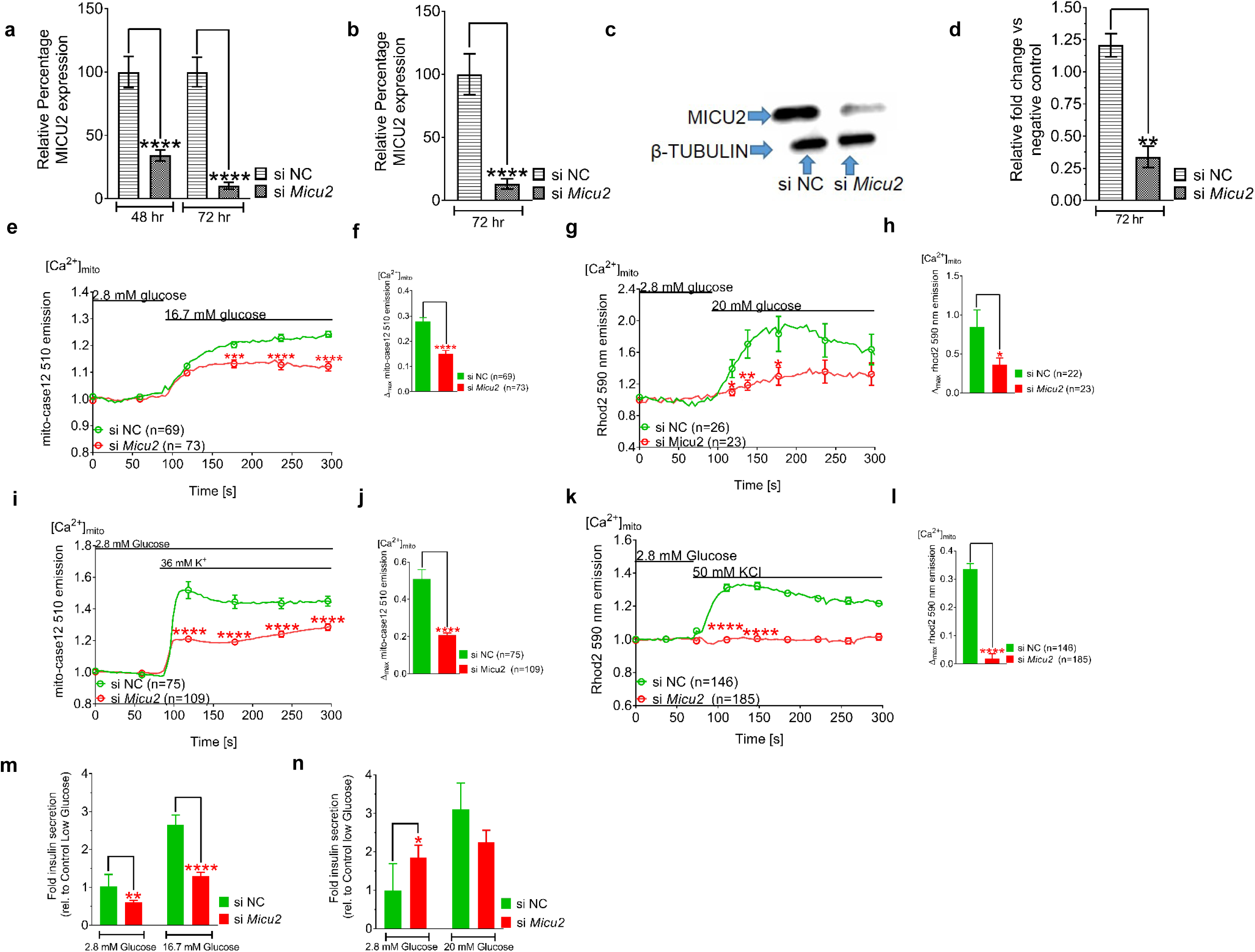
Knock down of *MICU2* reduces mitochondrial Ca^2+^ uptake and decreases insulin secretion. *MICU2* mRNA expression in INS-1 832/13 cells (A) and EndoC-βH1 cells (B) after treatment with scrambled or *MICU2* siRNA. MICU2 protein expression (C) and quantification of Western blots (D) in EndoC-βH1 cells. Mitochondrial Ca^2+^ ([Ca^2+^]_mito_) determined by mito-case12 and Rhod2, respectively, in *MICU2*-silenced INS-1 832/13 (E, F) and EndoC-βH1 cells (G, H) after glucose stimulation. E and G show average traces while F and H show differences in maximal mitochondrial Ca^2+^ levels between control and *MICU2*-silenced cells. [Ca^2+^]_mito_ by mito-case12 and Rhod2, respectively, in *MICU2*-silenced INS-1 832/13 (I, J) and EndoC-βH1 cells (K, L) after stimulation with KCl; average traces (I, K) and differences in maximal [Ca^2+^]_mito_ (J, L) between control and *MICU2*-silenced cells are shown. Insulin secretion in INS-1 832/13 (M) and EndoC-βH1 cells (N); secretion after a 1 h incubation was normalized to levels in cells treated with scrambled siRNA at 2.8 mM glucose. Data are from at least three independent experiments 72 h after addition of siRNA, unless otherwise stated. Mean ± SEM are given. Comparisons were made with an unpaired two-tailed Student’s t-test. * P<0.05; ** P<0.01; *** P<0.001; **** P<0.0001. See also Figure S1.

Given the proposed function of MICU2 in the uniporter holocomplex, we examined uptake of Ca^2+^ into the mitochondria upon *MICU2* knock down. To this end, we transfected INS-1 832/13 cells with a genetically encoded Ca^2+^ biosensor targeted to the mitochondrial matrix – mito-case12 (Zhdanov, Waters et al., 2014). Our experiments revealed that 16.7 mM glucose induced [Ca^2+^]_mito_ elevation and that this increase was attenuated by 48% after silencing of *MICU2* (*p* < 0.0001; Figure 1E-F). Similar findings were made in EndoC-βH1 cells using another reporter – Rhod2 – as these cells are difficult to transfect with plasmids (Logan, Szabadkai et al., 2014). In these cells, 20 mM glucose triggered a [Ca^2+^]_mito_ increase that was reduced by 57 % upon *MICU2*-silencing (*p* = 0.04; Figure 1 G-H). In addition, *MICU2* knock down reduced not only the magnitude but also the rate of [Ca^2+^]_mito_ elevation in EndoC-βH1 cells (113 s versus 65 s to maximal [Ca^2+^]_mito_; *p* = 0.0202; Supplementary Figure 1C). It should also be noted that both Rhod2 and mito-case12 showed ‘punctate’ loading, reflective of mitochondrial localization (Supplementary Figure 1D-E).

Raising extracellular K^+^ to 36 mM depolarized the plasma membrane and activated voltage-dependent Ca^2+^-channels. This resulted in a robust rise in [Ca^2+^]_mito_ in control cells, while there was only a minimal response in *MICU2*-silenced INS-1 832/13 cells (*p* < 0.0001; Figure 1I-J). Thus, the diminished mitochondrial response to elevated glucose (Figure 1, E-H) cannot simply be ascribed to a decreased coupling between glucose metabolism and plasma membrane depolarization. Again, EndoC-βH1 cells behaved similarly to the rodent insulinoma cells, where the maximal [Ca^2+^]_mito_ elevation following depolarization with 50 mM KCl was diminished by 90% in *MICU2*-silenced cells (*p* < 0.0001; Figure 1K-L).

Thus far, our data suggest that MICU2 plays a role in [Ca^2+^]_mito_ homeostasis in insulin-secreting cell lines, which would be predicted to impact hormone secretion. Indeed, we found that GSIS was reduced by 41% and 51% at 2.8 (*p* = 0.0023) and 16.7 mM glucose (*p* < 0.0001), respectively, in *MICU2*-deficient INS-1 832/13 cells (Figure 1M). In contrast, in *MICU2*-deficient EndoC-βH1 cells, basal insulin secretion at 2.8 mM glucose was increased by 46 % (*p* = 0.0245; Figure 1N). Secretion at 20 mM glucose, however, was not increased; accordingly, the net fold change (2.8 mM to 20 mM glucose) in insulin secretion was markedly and significantly reduced (1.4-fold vs. 3.1-fold; *p* = 0.0018).

### MICU2-Deficiency Impairs Bioenergetic Functions in Insulin-Secreting Cells

The mitochondrial membrane potential, Δψ_m_, is an essential facet of mitochondrial bioenergetics underlying both ATP synthesis and Ca^2+^-uptake into the mitochondrial matrix via MCU (Ward, Rego et al., 2000). The mitochondrial membrane potential is influenced by uptake of Ca^2+^ into the matrix. Given the potential regulatory role of MICU2 in this process and the observed attenuation of mitochondrial Ca^2+^ uptake, we monitored relative changes in Δψ_m_ with tetramethylrhodamine methylester (TMRM) in quench mode. Indeed, a rise in glucose from 2.8 to 16.7 mM hyperpolarized Δψ_m_ in INS-1 832/13 cells and this response was decreased by 50% after silencing of *MICU2* (*p* < 0.0001; Figure 2A-B). It therefore appears plausible that a decreased TCA cycle activation after silencing is the cause of the decreased hyperpolarization.

**Figure 2.**
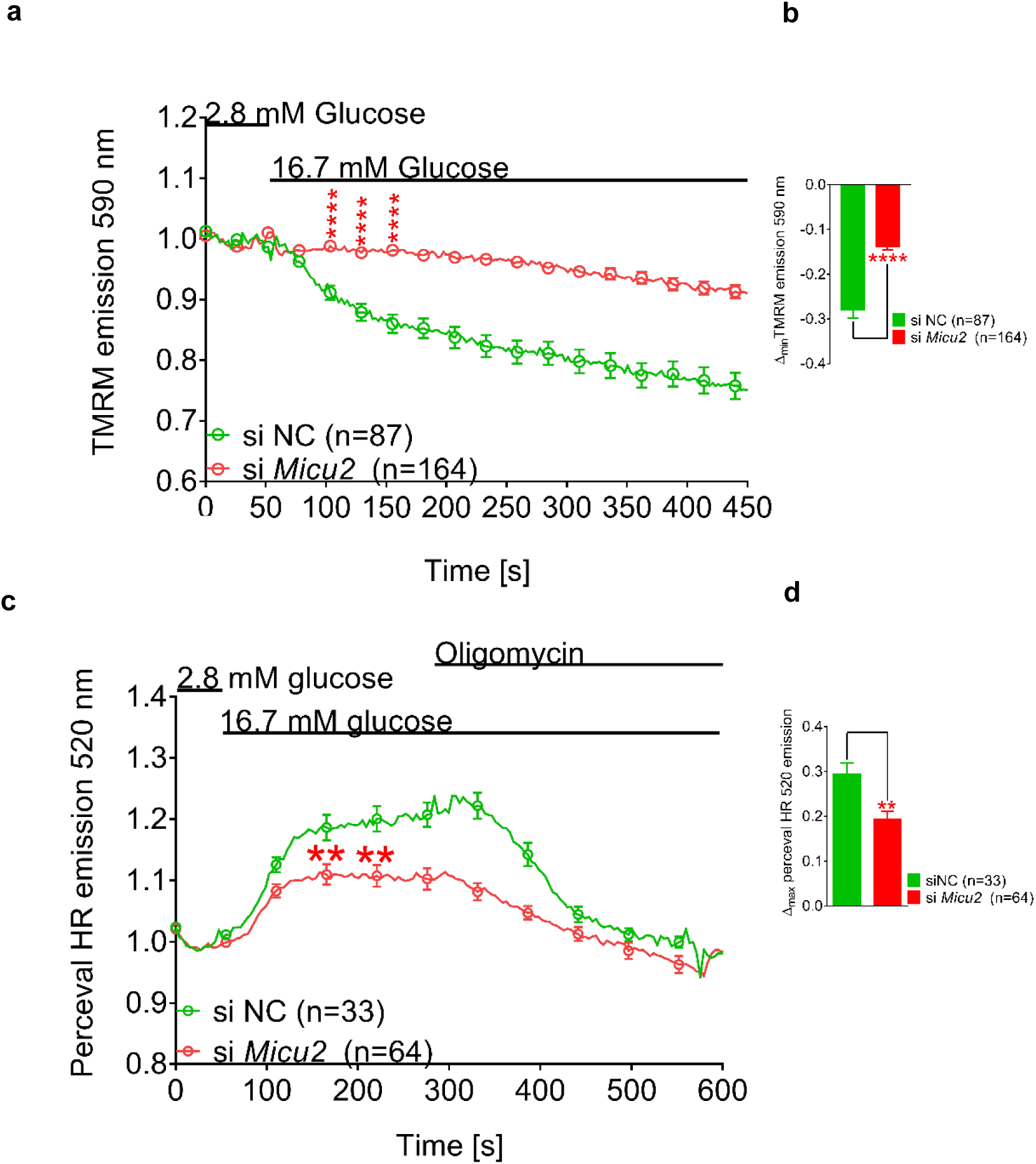
Bioenergetic consequences of *MICU2* knock down in INS-1 832/13 cells. Glucose-induced mitochondrial hyperpolarization of the inner mitochondrial membrane upon *MICU2* knock down in INS-1 832/13 cells measured by TMRM (A, B); average traces (A) and difference in maximal hyperpolarization (B) between control and *MICU2*-silenced cells are shown. Cytosolic ATP/ADP ratio in response to an elevation of glucose from 2.8 to 16.7 mM upon *MICU2* knock down in single INS-1 832/13 cells monitored with Perceval HR (C, D); average traces (C) and difference in maximal rise in ATP/ADP ratio (D) between control and *MICU2*-silenced cells are shown. Data are from at least three independent experiments 72 h after addition of siRNA. Mean ± SEM are given. Comparisons were made with an unpaired two-tailed Student’s t-test. ** P<0.01; **** P<0.0001.

In the light of this, we examined the effect of *MICU2*-deficiency on glucose-stimulated changes in the cytosolic ATP/ADP ratio, using live cell imaging of INS-1 832/13 cells expressing the fluorescent ATP/ADP reporter Perceval HR (Tantama, Martinez-Francois et al., 2013). We found that knock down of *MICU2* significantly reduced the rise in the cytosolic ATP/ADP ratio upon stimulation with 16.7 mM glucose (Figure 2C), with the maximal cytosolic ATP/ADP ratio increase diminished by 36 % after stimulation with 16.7 mM glucose in *MICU2*-silenced INS-1 832/13 cells (*p* = 0.0013; Figure 2C-D). However, as stated above, the decreased [Ca^2+^]_mito_ response to elevated glucose cannot be solely ascribed to decreased bioenergetic coupling, since it is also apparent when KCl was elevated.

### MICU2 Controls Both Mitochondrial and Cytosolic Ca^2+^ Entry

In the next set of experiments, we examined whether MICU2 also influences bulk [Ca^2+^]_c_ in insulin-secreting cells. We predicted that depolarization-evoked [Ca^2+^]_c_ increases would be more pronounced in *MICU2*-silenced cells due to reduced buffering by the mitochondria. Surprisingly, we observed that the maximal [Ca^2+^]_c_ rise upon stimulation with 36 mM KCl was reduced by 48 % in *MICU2*-silenced INS-1 832/13 cells (*p* < 0.0001; Figure 3A-B). Furthermore, the response was slower, with maximal [Ca^2+^]_c_ elevation reached in 50 seconds in *MICU2*-deficient cells compared to 21 seconds in the control (*p* < 0.0001; Supplementary Figure 2A). Similar results were obtained in EndoC-βH1 cells, where silencing of *MICU2* led to a 41 % reduction in the maximal [Ca^2+^]_c_ elevation upon 50 mM KCl stimulation (*p* = 0.0019; Supplementary Figure 2B-C). This implies that the role of MICU2 in this paradoxical regulation of [Ca^2+^]_c_ is conserved in β-cells of both rat and human origin.

**Figure 3.**
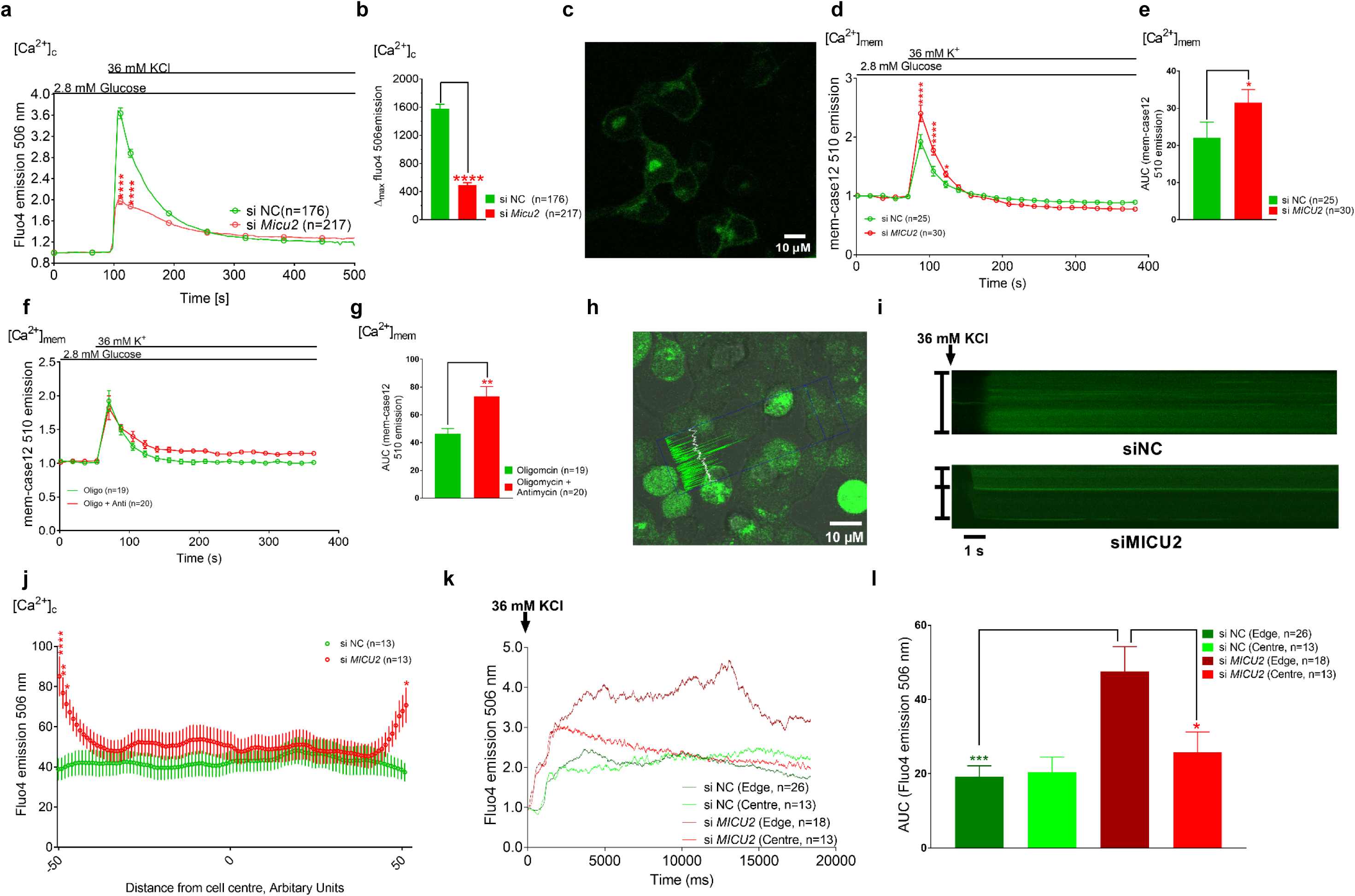
Cytosolic and sub-plasmalemmal Ca^2+^ levels in response to KCl stimulation is perturbed in *MICU2*-silenced INS-1 832/13 cells. Cytosolic Ca^2+^levels ([Ca2+]_c_) in MICU2-silenced INS-1 832/13 cells (and scramble control) upon stimulation with 36 mM KCl, measured by Fluo4 (A, B); Average traces are shown in A, while B shows differences in maximal Fluo4 signals. (C) Image of mem-Case12 staining, n.b. specific membrane localisation. Submembrane Ca^2+^ levels ([Ca^2+^]_mem_ in *MICU2*-silenced INS1 832/13 cells upon stimulation with 36 mM KCl, measured using mem-case12 (D, E); Average traces are shown in D, while E shows iAUC quantification of KCl effect on [Ca^2+^]_mem_. In (F, G) similar experiments are shown but using oligomycin and antimycin to block mitochondrial Ca^2+^ uptake prior to KCl addition, with oligomycin applied alone as a control. Mean ± SEM are given. (H) Example of linescan selection: the blue arrow going through the centre of the cell is the only part of the field of view that is imaged; the accompanying trace within the blue rectangle shows the fluorescence signal for each corresponding point along the aforementioned arrow. (I) Representative linescan recordings of cytosolic Ca^2+^ levels in MICU2-silenced INS1 832/13 cells (and scramble control) in response to 36 mM KCl, measured by Fluo4. Double-headed flat-headed arrows represent each individual cell’s axis (which is where the Ca^2+^ signal is measured); note the higher Ca^2+^ signal at the membrane in siMICU2-treated cells, reflected by the thin lines of higher intensity at the edge of each cell. (J) Mean Ca^2+^ intensity across INS1-832/13 cells, ∼500 ms after initiation of response to 36 mM KCl. Cells were of different sizes, therefore data were interpolated to standardize each cell to a set number of points, allowing for comparison. The X-axis represents cell axis, moving from one side of the plasma membrane (−50) through the centre of the cell (0) to the opposite side of the plasma membrane (50). Points close to the extremities represent the submembrane region. (K) Average trace of cytosolic Ca^2+^ at cell edge and cell centre in *MICU2*-silenced INS1 832/13 cells (and scramble control). (L) iAUC quantification of [Ca^2+^]_c_ at cell edge and cell centre in *MICU2*-silenced INS1 832/13 cells (and scramble control). Sub-plasmalemmal Ca^2+^ levels ([Ca^2+^]_mem_) in *MICU2*-silenced INS-1 832/13 cells (and scramble control) upon stimulation with 36 mM KCl. Comparisons were made with an unpaired two-tailed Student’s t-test. *P<0.05; ** P<0.01; ***P<0.001; **** P<0.0001.

We hypothesized that the reduced depolarization-evoked [Ca^2+^]_c_ increases after *MICU2* knock down (and mitochondrial depolarization) were due to reduced net Ca^2+^ entry across the plasma membrane (diminished entry or enhanced extrusion). To elucidate this functionally, we imaged INS1-832/13 cells, using a genetically encoded Ca^2+^ biosensor targeted to the inner leaflet of the plasma membrane – mem-case12 (Nagai, Sawano et al., 2001, Skene & Virag, 1989, Souslova, Belousov et al., 2007). Figure 3C shows the plasma membrane localization of the sensor, allowing the sub-plasmalemmal free Ca^2+^ concentration ([Ca^2+^]_mem_) to be determined. If mitochondrial Ca^2+^ sequestration was indeed facilitating net Ca^2+^ entry into the cell, by removal of Ca^2+^ from the sub-plasma membrane compartment, then it would be predicted that cells with defective mitochondrial Ca^2+^ transport would show enhanced sub-plasmalemmal Ca^2+^ elevation in response to depolarization by KCl. Indeed, the maximal [Ca^2+^]_mem_ elevation was 20% greater in *MICU2*-deficient INS-1 832/13 cells than in control cells (*p* = 0.0118; Figure 3H). The iAUC was also 43% greater in the *MICU2*-deficient cells (*p* = 0.015 vs. control; Figure 3I).

To establish whether this effect on [Ca^2+^]_mem_ could be accounted for by restriction of MCU activity, we used an alternative strategy to inhibit mitochondrial Ca^2+^ transport (Budd & Nicholls, 1996). In the presence of the ATP synthase inhibitor, oligomycin, ATP is maintained by glycolysis and Δψ_m_ is retained, or even elevated, allowing mitochondrial Ca^2+^ transport to occur normally. The further addition of an electron transport inhibitor, such as antimycin, rapidly collapses Δψ_m_ with no a priori effect on (glycolysis-maintained) cytosolic ATP production. Mitochondrial Ca^2+^ transport, however, being dependent upon Δψ_m_, is abolished (Budd & Nicholls, 1996). We found that although treating control INS-1 832/13 cells with oligomycin/antimycin did not increase the maximal [Ca^2+^]_mem_ elevation, the Ca^2+^ concentration remained elevated at the membrane for longer with a delayed return to the basal state (Figure 3J). This was reflected by the iAUC of the effect increasing by 58 % compared to oligomycin control (*p* = 0.0026; Figure 3K). Thus, this treatment had a comparable effect on [Ca^2+^]_mem_ as did *MICU2*-deficiency. An important proviso is that the capacity of INS-1 832/13 cells to accelerate glycolysis in the presence of oligomycin is limited by their low expression of lactate dehydrogenase (Malmgren, Nicholls et al., 2009); therefore, experiments were performed during the time line before which ATP depletion became significant, as indicated by the unperturbed mitochondrial membrane potential during this time (data not shown). Furthermore, using oligomycin alone as a control implies that any effect upon ATP production cannot explain the reduced extrusion of [Ca^2+^]_mem_ in oligomycin/antimycin-treated INS1 832/13 cells and must therefore be due to a decrease in mitochondrial Ca^2+^ uptake.

To further examine whether sub-plasmalemmal Ca^2+^ elevations occur when mitochondrial Ca^2+^ uptake is abrogated, we imaged INS1-832/13 cells loaded with Fluo4, using the linescan configuration. This allows repeated imaging across a line through a cell(s) axis as opposed to the entire field of view (Figure 3H). Crucially, this method allows for the higher temporal resolution required for tracking Ca^2+^ signal propagation following membrane depolarization. To this end, cell(s) were selected, after which the starvation medium was rapidly replaced with medium containing 36 mM KCl and the recording started simultaneously. It was expected that MICU2-deficient cells would show enhanced sub-plasmalemmal Ca^2+^ elevation in response to depolarization by KCl, reflected by a stronger fluorescence signal intensity at the edge of the linescan. Indeed, we found that, upon KCl-induced depolarization, Ca^2+^ was accumulating at the sub-plasma membrane compartment in si*MICU2*-treated INS1-832/13 cells (Figure 3I-K) with a 1.85-fold higher Ca^2+^ signal (measured as incremental area under the curve (iAUC)) at the edge of the cell compared to its centre (*p* = 0.0266). Furthermore, there was a 2.5-fold higher Ca^2+^ response at the submembrane compartment in *MICU2*-deficient INS1-832/13 cells compared to control (*p* = 0.0006 vs. control; Figure 3L). In contrast, there was no significant difference in the Ca^2+^ signal between the edge and centre in control cells (*p* = 0. 8027; Figure 3L). Note that when analyzing the linescan data the nucleus was avoided when selecting the ‘centre’ of the cell.

Collectively, these observations support a model where *MICU2*-deficient mitochondria in the vicinity of the plasma membrane are less capable of removing Ca^2+^ from this compartment. Our results thus indicate not only that MICU2 plays a critical role in the control of the MCU and Ca^2+^ homeostasis, but also that mitochondria play a critical role in buffering local sub-plasmalemmal Ca^2+^, facilitating net Ca^2+^ entry into the cell. This may occur either by decreasing cytosolic Ca^2+^-mediated desensitization of voltage-dependent Ca^2+^ channels in the plasma membrane (DeCaen, Liu et al., 2016, Navedo, Amberg et al., 2005), or alternatively by restricting Ca^2+^ efflux by plasma membrane Ca^2+^-ATPases (Demaurex, Poburko et al., 2009, Groten, Rebane et al., 2016). The latter is unlikely in the present experimental situation since *MICU2*-deficiency inhibited a rise in the ATP/ADP ratio, which will limit access to ATP for the ATPases.

### *Micu2*-deficient Mice Show Defective Stimulus-Secretion Coupling and Insulin Secretion

Our experiments until now suggested that perturbation of MICU2 leads to reduced mitochondrial Ca^2^ uptake, resulting in perturbed Ca^2+^ entry into the cell and, consequently, defective insulin secretion in both rodent derived INS-1 832/13 and human EndoC-βH1 cell lines. To understand the consequences of MICU2-deficiency in a more physiological setting, we used a global *Micu2* knock out mouse (Bick, Wakimoto et al., 2017). As previously observed in the insulin-secreting cell lines, glucose-stimulated (20 mM) maximal [Ca^2+^]_mito_ elevation was diminished by 57 % in *Micu2* knock out islets as compared to wild type islets (*p* < 0.0001; Figure 4A-B). Note that the KCl-response was also diminished (Figure 4A), similar to that observed in the cell lines. These results thus suggest that MICU2 most likely plays a similar role in maintenance of [Ca^2+^]_mito_ homeostasis in primary cells to that observed in cell lines.

**Figure 4.**
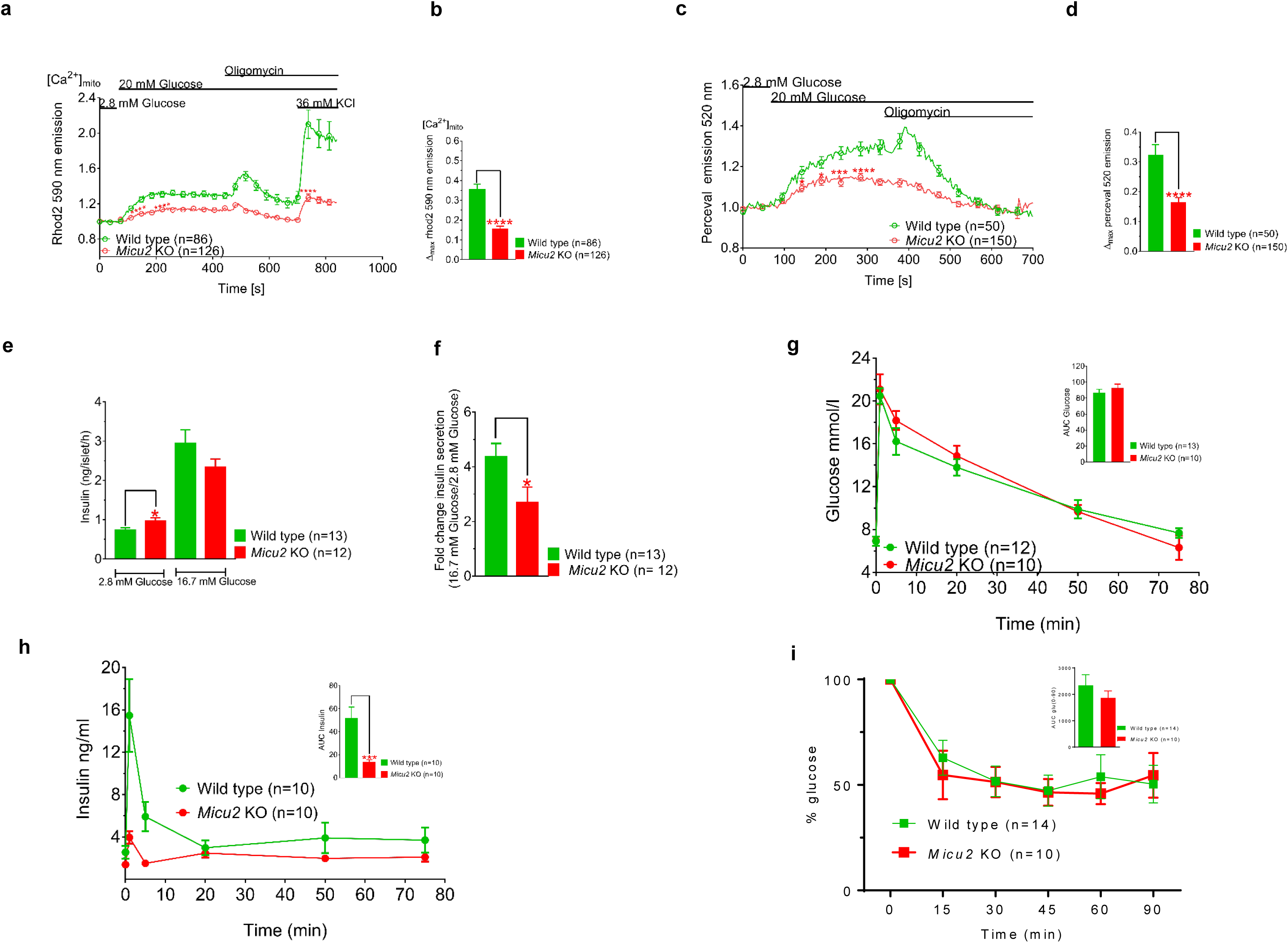
*MICU2* knock out mice exhibit deficiencies in mitochondrial Ca^2+^ homeostasis, ATP/ADP ratio, and insulin secretion. Mitochondrial Ca^2+^ levels [Ca^2+^]_mito_ in islets isolated from *Micu2* knock out (KO) and wild type mice measured with Rhod2 after stimulation with 20 mM glucose (A, B). Average traces are shown in (A) while (B) show differences in maximal Rhod2 signals. 40 islets from 7 wild type mice and 42 islets from 6 *Micu2* KO mice were assayed in response to 20 mM glucose and 36 mM KCl. Changes in the cytosolic ATP/ADP ratio in response to 20 mM glucose in *Micu2* and wild type islets were determined by Perceval (C); n=14 and 27 islets from 4 and 6 wild type and KO islets, respectively. Average traces are shown in (C) while (D) show differences in maximal Perceval signals. Insulin secretion from *Micu2* KO and wild type mice at 2.8 and 16.7 mM glucose in a 1 h incubation (E); (F) shows fold-response of insulin. Plasma glucose (G) and insulin (H) levels in an intravenous glucose tolerance test in *Micu2* KO and wild type mice; n denotes number of mice. Insets show area under curve for glucose (AUC; H) and insulin (J) T=0 – 5 minutes. Data are from at least three independent experiments. (I) Insulin tolerance test (ITT) in *Micu2* KO and wild type mice. Glucose values are expressed as % glucose where glucose at 0 min is 100%; n denotes number of mice. Inset shows AUC. Mean ± SEM are given. Comparisons for imaging were made with an unpaired two-tailed Student’s t-test (**** P<0.0001.) and Mann-Whitney U-test (***P<0.001) for data from IVGTT.

Furthermore, as [Ca^2+^]_mito_ homeostasis is critical for cellular bioenergetics, we speculated that the *Micu2* knock out might perturb the cytosolic ATP/ADP ratio in primary cells as was seen in the insulin-secreting cell lines. Therefore, in the next set of experiments, we used an adenovirus expressing the biosensor Perceval, originally described as an ATP/ADP sensor (Berg, Hung et al., 2009), but later shown to report cytoplasmic ATP in mouse islet cells (Li, Yu et al., 2015). The maximal Perceval signal was diminished by 50 % in *Micu2* knock out islets compared with wild type islets upon stimulation with 20 mM glucose. (*p* < 0.0001; Figure 4C-D). A likely consequence of an abrogated rise in the ATP/ADP ratio in pancreatic β-cells is impaired insulin secretion. Batch incubations of *Micu2* knock out islets demonstrated that basal (2.8 mM glucose) insulin secretion was significantly higher as compared to that in their wild type counterparts (*p* = 0.0427; Figure 4E). However, *Micu2* knock out islets failed to release insulin in response to high glucose as efficiently as wild type islets, illustrated by a 50 % reduction in the fold response of insulin secretion (16.7 to 2.8 mM glucose; *p* = 0.0362; Figure 4F). Our experiments thus suggest that stimulus-secretion coupling in β-cells in *Micu2* knock out mice is defective and underlies perturbed GSIS, reminiscent of that found in EndoC-βH1 cells (Figure 1N).

In the next set of experiments, we examined the consequences of *Micu2*-deficiency on whole body glucose homeostasis. Glucose elimination was not significantly different between the *Micu2* knock out and wild type mice (Figure 4G). However, an intravenous glucose tolerance test (IVGTT) revealed a marked reduction of the initial release of insulin (*p* = 0.0003; Figure 4H): we observed a 74 % and 70 % reduction in the plasma insulin levels at 1 and 5 minutes after glucose administration, respectively, in *Micu2* knock out mice compared to wild type mice. In view of that insulin secretion was impaired while glucose elimination was unaltered in *Micu2*-deficiency, we performed an insulin tolerance test (ITT). Both strains of mice responded robustly to insulin with lowering of plasma glucose levels (Figure 4I). These experiments thus demonstrate that the initial release of insulin in *Micu2* knock out mice was reduced, likely caused by a stimulus-secretion coupling defect in pancreatic β-cells. Hence, *Micu2* knock out mice exhibited deficient GSIS both *in vitro* and *in vivo*.

### MICU2-regulation of Ca^2+^ in Mitochondria, Cytosol and Sub-Plasmalemmal Space is Conserved Between Cell Types

Previous studies in neuronal cells have shown that inhibition of ATP generation and dissipation of mitochondrial membrane potential lead to reduced [Ca^2+^]_c_ upon stimulation with glutamate or 50 mM KCl (Castilho, Hansson et al., 1998). Therefore, having shown the role of MICU2 in regulation of [Ca^2+^]_mito_ and [Ca^2+^]_c_ in insulin-secreting cells, we asked whether this mechanism may be observed also in other cell types. High throughput gene expression assays have previously demonstrated that the human embryonic kidney-derived HEK-293T cell line expresses robust levels of *MICU2* mRNA (Zolg, Wilhelm et al., 2017). Furthermore, it has been shown that this cell line also expresses voltage-dependent Ca^2+^ channels and exhibits endogenous Ca^2+^ currents (Berjukow, Doring et al., 1996, Wang, Davis et al., 2005). We therefore used this cell line to address the role of MICU2 in cellular Ca^2+^ homeostasis.

*MICU2* was effectively silenced on the mRNA and protein levels in HEK-293T cells (Supplementary Figure 3A-B). Measurements of [Ca^2+^]_mito_ with mitocase-12 demonstrated that stimulation with 36 mM KCl increased [Ca^2+^]_mito_ in HEK293T cells and that this response was attenuated by 46% after silencing of *MICU2* (*p* < 0.0001; Figure 5A-B). In addition, time to maximal [Ca^2+^]_mito_ peak in *MICU2*-silenced cells was 126 seconds as compared 107 seconds in controls after 36 mM KCl stimulation (*p* = 0.0087; Supplementary Figure 3D). [Ca^2+^]_c_ measurements using Fluo5F showed that membrane depolarization by 50 mM KCl triggered a [Ca^2+^]_c_ increase and the maximal [Ca^2+^]_c_ elevation was reduced by 74 % in *MICU2*-silenced HEK-293T cells compared to control cells (*p* < 0.0001; Figure 5C-D). For experiments measuring [Ca^2+^]_mem_, using mem-case12, oligomycin was added to ensure that any effects of *MICU2* silencing were independent from the resultant bioenergetic effects of increased [Ca^2+^]_mito_. The same was not done in the experiments with INS1 832/13 cells due to ATP production in the β-cell being finely tuned to the extracellular glucose concentration as a consequence of the low affinity enzymes GLUT2 and glucokinase, and downstream dissipative pathways (i.e. high leak current) (Nicholls, 2016). This means that at low glucose, metabolic flux and ATP production in INS1 832/13 cells are low compared to HEK293T cells, therefore, any effect of increased [Ca^2+^]_mito_ on ATP production would be unlikely to reach the threshold to affect K_ATP_ channel activity, meaning use of oligomycin was not necessary. Reminiscent to the findings in INS1 832/13 cells, [Ca^2+^]_mem_ elevation upon 50 mM KCl stimulation was increased by 35 % in *MICU2*-silenced HEK293T cells as compared to the control cells (*p* = 0.0103; Figure 5E-F). Thus, these data demonstrate a conservation of the regulation of [Ca^2+^]_mito_ and [Ca^2+^]_cyto_ by MICU2 in different cell types.

**Figure 5.**
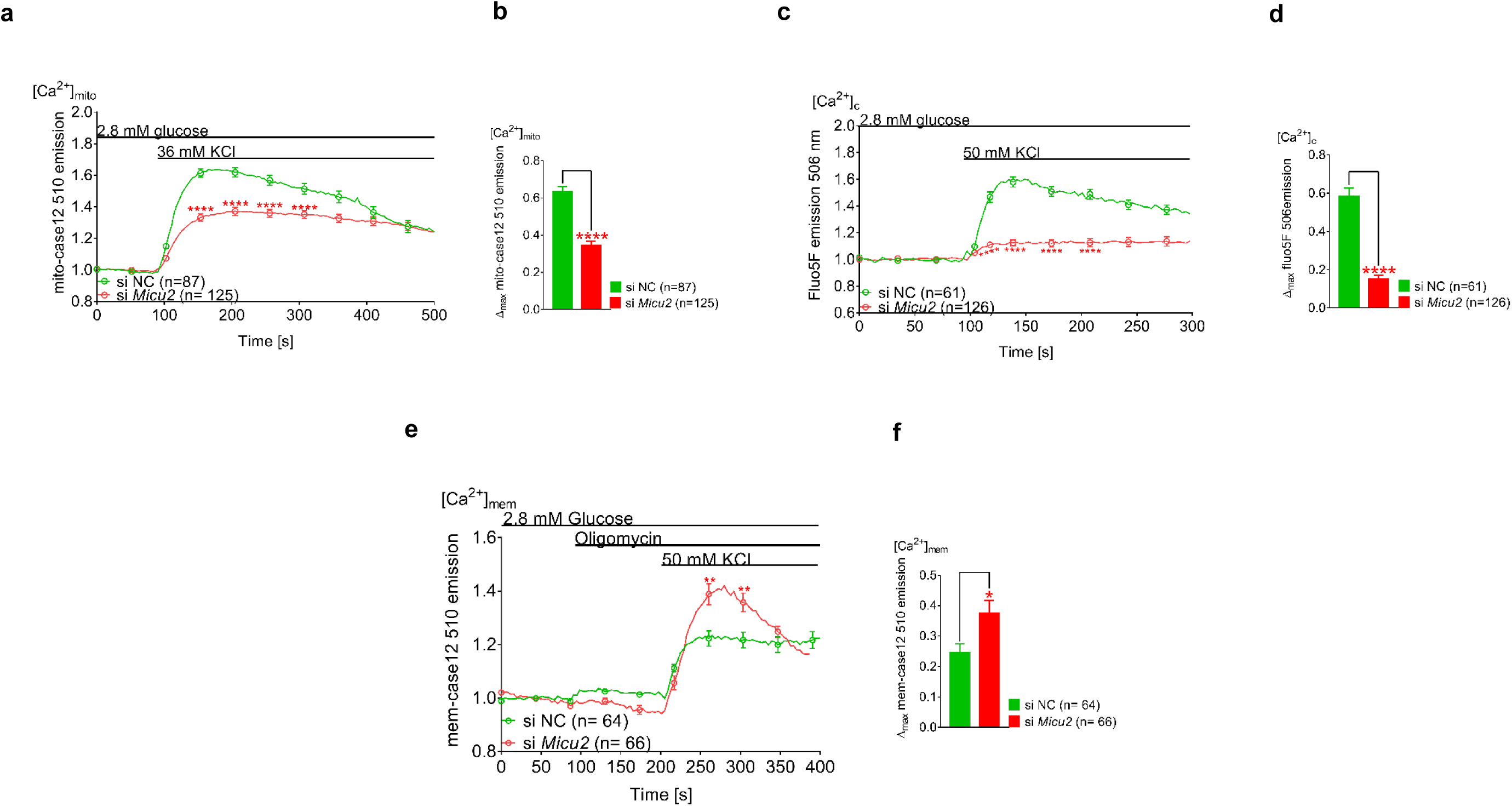
MICU2 plays a similar role in Ca^2+^ homeostasis in HEK-293T cells as observed in insulin-secreting cells. Mitochondrial Ca^2+^ ([Ca^2+^]_mito_) determined by mito-case12 in *MICU2*-silenced HEK-293T cells stimulated with 36 mM KCl (A, B). (A) shows average traces while (B) shows differences in maximal mitochondrial Ca^2+^ response. Cytosolic Ca^2+^ levels ([Ca^2+^]_c_) determined by Fluo5F in *MICU2*-silenced HEK-293T cells upon stimulation with 50 mM KCl (C, D); Average traces are shown in (C) while (D) shows differences in maximal Fluo5F signals. Sub-plasmalemmal Ca^2+^ levels ([Ca^2+^]_mem_) in *MICU2*-silenced HEK-293T cells upon stimulation with 50 mM KCl (E, F); average traces are shown in (E) while (F) shows differences in maximal mem-case12 signals. Data are from at least three independent experiments 72 h after addition of siRNA. Mean ± SEM are given. Comparisons were made with an unpaired two-tailed Student’s t-test. * P<0.05; ** P<0.01; **** P<0.0001. See also Figure S3.

## DISCUSSION

It is undisputed that a rise in [Ca^2+^]_c_ triggers insulin secretion, as well as secretion of most other hormones, but it has also been recognized that such a rise in itself does not sustain insulin secretion (Gembal, Detimary et al., 1993, Jonas, Li et al., 1994). Cytosolic Ca^2+^ is further transported into the mitochondrial matrix, and an elevation in free matrix Ca^2+^ has been considered to trigger additional regulatory factors (Rutter, Theler et al., 1993). Prior to the molecular identification of the components of the uniporter holocomplex (Baughman, Perocchi et al., 2011, Perocchi, Gohil et al., 2010, Plovanich, Bogorad et al., 2013), indirect approaches were required to investigate the effect of modulating [Ca^2+^]_mito_, none of which were entirely satisfactory. Removal of external Ca^2+^ (Balke, Egan et al., 1994) affects multiple non-mitochondrial processes, not least exocytosis itself. Moreover, attempts to dampen changes in [Ca^2+^]_mito_ by increasing the matrix Ca^2+^ buffering capacity, by targeting exogenous Ca^2+^-binding proteins to the matrix (Wiederkehr, Szanda et al., 2011) or by loading cells with the Ca^2+^-chelator 1,2-bis(o-aminophenoxy)ethane-*N,N,Ń,Ń-*tetraacetic acid (BAPTA) (Tanaka, Nagashima et al., 2014), each succeeded in reducing the bioenergetic response to elevated glucose, but concerns remain about toxicity associated with the treatment (Nicholls, 2016).

The molecular characterization of the uniporter (Baughman et al., 2011, De Stefani, Raffaello et al., 2011, Perocchi et al., 2010, Plovanich et al., 2013), as well as a number of accessory proteins, i.e., MICU1, 2, 3 and EMRE (Perocchi et al., 2010, Plovanich et al., 2013, Sancak et al., 2013) opened the possibility of genetic manipulation of the components of the uniporter (Jhun et al., 2016). It was shown that silencing of *Mcu* abrogates the rise in [Ca^2+^]_mito_ and the cytosolic ATP/ADP ratio in response to glucose in mouse β-cells (Tarasov, Semplici et al., 2012). Another study demonstrated that silencing of *MCU* in clonal INS1-E cells results in reduced nutrient-induced hyperpolarization of the inner mitochondrial membrane, ATP production and GSIS (Quan, Nguyen et al., 2015). Moreover, knock down of *MICU1* leads to reductions in [Ca^2+^]_mito_, cytosolic ATP levels and GSIS in clonal insulin-secreting cells (Alam et al., 2012). The role of MICU2, however, remains more enigmatic. MICU2 is a peripheral membrane protein associated with the mitochondrial inner membrane and facing the intermembrane space. The absence of MICU2 has been associated with reduced Ca^2+^ uptake *in vivo* in mouse liver, though silencing in this system also leads to a decrease in the abundance of the pore-forming subunit MCU and the complex (Plovanich et al., 2013). Observations in other cell systems suggest that the accessory protein may have a regulatory role in Ca^2+^ uptake into mitochondria (Kamer & Mootha, 2014, Patron, Checchetto et al., 2014). MICU1 and MICU2 are thought to heterodimerize, by means of disulfide bridges, and thus regulate Ca^2+^ entry into the mitochondrial matrix via the MCU (Tsai, Wu et al., 2017). Indeed, both MICU1 and MICU2 have been shown to play an important role in setting the threshold for mitochondrial Ca^2+^ uptake in HeLa cells at variable Ca^2+^ concentrations (Patron et al., 2014). Furthermore, loss of MICU1 or MICU2 in permeabilized HEK-293T cells has also been shown to disrupt this critical threshold for mitochondrial Ca^2+^ uptake (Kamer et al., 2017, Kamer, Jiang et al., 2019, Kamer & Mootha, 2014, Xing, Wang et al., 2019).

Here, through use of *MICU2*-deficient rodent and human insulin-producing cell lines, as well as *Micu2* knock out mice, we were able to show that reduced Ca^2+^ uptake into mitochondria resulted in inhibition of GSIS. This confirmed observations in previous studies (Alam et al., 2012, Quan et al., 2015, Tarasov et al., 2012) showing that [Ca^2+^]_mito_ affects mitochondrial metabolism and insulin secretion. Thus far, our experiments could not resolve how *Micu2* knock out mice maintained euglycemia despite reduced early insulin secretion. Insulin tolerance, albeit a rough estimate of insulin action *in vivo*, was unchanged. In fact, we have previously reported that ITT is not sensitive enough to reflect changes in insulin sensitivity in the face of reduced early insulin secretion *in vivo* (Fex, Haemmerle et al., 2009). A hyperinsulinemic euglycemic clamp, which is beyond the scope of the present studies, would be required to accurately determine insulin sensitivity. Importantly, *MCU* and *MICU1* expression were unaffected by *MICU2* silencing in INS1 832/13 cells, indicating that MICU2 plays a fundamental, non-redundant, role in regulating [Ca^2+^]_mito_ in β-cells. However, the mechanisms by which MICU2 and [Ca^2+^]_mito_ might exert control over GSIS were unexpected and add a new dimension to how mitochondrial Ca^2+^ transport might regulate β-cell function and insulin secretion.

The key observation, novel in the context of the β-cell, is that mitochondrial Ca^2+^ uptake exerts a profound stimulatory effect on the net flux of the cation across the plasma membrane, either by enhancing entry or depressing efflux. Insulin-secreting cells have a high concentration of sub-plasma membrane mitochondria, with roughly 20% of the membrane inner surface associated with mitochondria exerting a strong Ca^2+^ buffering effect (Griesche, Sanchez et al., 2019). Mitochondria are dynamic organelles and sub-membrane Ca^2+^ buffering is reduced upon plasma membrane depolarization because of Ca^2+^-dependent remodeling of the cortical F-actin network, causing translocation of sub-plasmalemmal mitochondria to the cell interior (Griesche et al., 2019). In line with these findings, we demonstrate that restricted mitochondrial Ca^2+^ uptake resulted in more pronounced depolarization-induced increases of Ca^2+^ in the sub-membrane compartment. However, under the same conditions, the global cytoplasmic Ca^2+^ increases were attenuated.

In order to rationalize a decreased Ca^2+^-response to plasma membrane depolarization in both the matrix and cytosolic compartments, the simplest assumption is that net Ca^2+^ influx across the plasma membrane is inhibited. Increased extrusion of Ca^2+^ is less likely, given the observed lowering of the ATP/ADP ratio as a consequence of MICU2-deficiency and that extrusion is an ATP-dependent process. While the endoplasmic reticulum does take up Ca^2+^ in response to the elevated ATP/ADP ratio at high glucose, the response is too slow to account for a Ca^2+^-deficit (Ravier, Daro et al., 2011). In any case, the effect of restricted mitochondrial Ca^2+^ uptake on bulk [Ca^2+^]_c_ is seen with elevated KCl, as well as with high glucose, and is thus not directly dependent on bioenergetics. Had this effect only been observed in response to glucose, then the previously postulated positive effect of mitochondrial Ca^2+^ uptake on mitochondrial dehydrogenases would have been the most plausible (Wiederkehr & Wollheim, 2008).

A paradoxical decrease in [Ca^2+^]_c_ elevation following inhibited transport of mitochondrial Ca^2+^ has previously been observed in cultured neurons (Budd & Nicholls, 1996, Castilho et al., 1998). Since these studies predated the molecular identification of the MCU, mitochondrial Ca^2+^-transport was inhibited by mitochondrial depolarization in the presence of the ATP synthase inhibitor oligomycin together with an electron transport inhibitor – rotenone or antimycin. The control was oligomycin alone: both control and mitochondrial depolarized cells maintain similar ATP/ADP ratios, supported by glycolysis, but the [Ca^2+^]_c_ transients induced by Ca^2+^ entry through NMDA-selective glutamate receptors or voltage-dependent Ca^2+^ channels are drastically reduced. In the present study, the treatment with oligomycin + antimycin further confirmed a role of mitochondrial Ca^2+^ uptake in clearing sub-plasmalemmal Ca^2+^ in insulin-secreting cells, and was similar to that found in neurons (Budd & Nicholls, 1996, Castilho et al., 1998). Importantly, the net neuronal accumulation of ^45^Ca^2+^ is strongly inhibited by mitochondrial depolarization (Budd & Nicholls, 1996, Castilho et al., 1998), confirming the ability of the organelle to control net flux across the plasma membrane. Notably, mitochondrial Ca^2+^ uptake has also been implicated in store-operated calcium entry with MCU knockdown in RBL mast cells shown to reduce Ca^2+^ influx through calcium-release activated channels (Samanta, Bakowski et al., 2018).

It was hypothesized that mitochondria are able to deplete a sub-plasma membrane layer of elevated Ca^2+^ (Budd & Nicholls, 1996, Castilho et al., 1998). Voltage-dependent Ca^2+^-channels become desensitized when local Ca^2+^ accumulations are not promptly removed (Demaurex et al., 2009, Schiff, Siderovski et al., 2000, Taylor, Huang et al., 2005). In addition, it has been shown in chromaffin cells, neurons and cardiac muscle that mitochondrial sequestration of sub-plasmalemmal Ca^2+^ prevents desensitization of Ca^2+^ channels and might thus uphold hormone secretion (David & Barrett, 2003, Montero, Alonso et al., 2000, Sanchez, Garcia et al., 2001). Furthermore, mitochondria play an essential role in maintenance of the [Ca^2+^]_c_ amplitude wave propagation via feedback regulation of plasma membrane channel activity in diverse cellular systems, including rat cortical astrocytes and *Xenopus leavis* oocytes (Boitier, Rea et al., 1999, Jouaville, Ichas et al., 1995). The response of the sub-plasmalemmal-targeted Ca^2+^ probe mem-case12 and linescan imaging in the present study provided support for the existence of such a layer of elevated Ca^2+^ (Budd & Nicholls, 1996, Castilho et al., 1998) during β-cell depolarization, and its depletion by mitochondrial Ca^2+^ uptake.

Notably, in the mem-case 12 experiments, there was a difference between the effects on [Ca^2+^]_mem_ between *Micu2* silencing and treatment with oligomycin/antimycin, whereby the former caused an increase in the duration of the subplasmalemmal Ca^2+^ increase while the latter resulted in an increased magnitude of the response. This may be explained by the different mechanisms by which mitochondrial Ca^2+^ uptake is inhibited. When MICU2 is deficient, uptake at low Ca^2+^ is likely to be increased as the protein plays an important role in suppressing MCU activity under these conditions (Kamer & Mootha, 2014, Matesanz-Isabel, Arias-del-Val et al., 2016, Patron et al., 2014). Subsequently, when [Ca^2+^]_c_ is elevated the buffering capacity of the mitochondria is reduced. In contrast, oligomycin and antimycin would prevent proton transport from the mitochondrial matrix to the mitochondrial intermembrane space, causing an increase in the matrix pH, subsequent collapsing of Dy_m_ and resultant inhibition of Ca^2+^ entry. These two methods would affect the Ca^2+^ electrochemical gradient differently and as a result the Ca^2+^ uptake kinetics, potentially explaining the differential effects on [Ca^2+^]_mem_.

Regarding the linescan imaging, one could argue that due to its relatively low K_d_, Fluo4 is unsuitable for measurement of [Ca^2+^]_mem_, which can reach concentrations of 10-100 µM (Parekh, 2008). This means there is a risk of Fluo4 saturating at the submembrane compartment. This did not seem to occur in the control, where there was little difference in the Fluo4 signal between the edge and centre of the cell. In *MICU2*-silenced cells, however, there was a clear difference between the edge and the centre of the cell. If saturation was indeed occurring, this would imply that the observed increase in [Ca^2+^]_mem_ was in fact an underestimation. Moreover, the problems of saturation were mitigated against with the complementary findings using the lower affinity mem-case12 probe (K_d_ = 1μM) (Souslova et al., 2007)..

Our present work shows that mitochondrial Ca^2+^ transport has additional consequences for the β-cell beyond facilitation of the TCA cycle, which may have varying effects on insulin secretion. On the one hand, attenuation of sub-plasmalemmal Ca^2+^ elevations could reduce the triggering of exocytosis of pre-docked vesicles. On the other hand, there is evidence that increased Ca^2+^ in the cytosol is required for the transport of reserve vesicles to the release site, at least in neuronal systems (Koenig, Yamaoka et al., 1993). In the present study, insulin secretion was determined during 1 h, i.e., mainly ‘second phase’ release, during which such mobilization of reserve vesicles may be quantitatively dominant.

Our results may help to resolve some of the contradictions in the concept of amplification pathways and K_ATP_-independent mechanisms of insulin secretion. The search for coupling factors responsible for K_ATP_-independent insulin secretion began after the discovery that insulin secretion could be provoked by increasing glucose even when the K_ATP_-channel was kept open by diazoxide and the plasma membrane was depolarized by high concentrations of extracellular KCl, thereby opening voltage-dependent Ca^2+^-channels (Gembal et al., 1993). Metabolic coupling factors were postulated to account for the insulinotropic effects of glucose under K_ATP_-independent conditions, as well as explain why insulin secretion could not be sustained by an elevation of [Ca^2+^]_c_ alone (Jonas et al., 1994). However, at least the latter circumstance can be accounted for by the consequences of mitochondrial Ca^2+^ uptake that we observed here: if Ca^2+^ is not removed from the sub-plasmalemmal space by mitochondria, voltage-dependent Ca^2+^-channels will be desensitized. A number of metabolites and metabolic pathways have been convincingly implicated in amplification of GSIS (Gooding, Jensen et al., 2016). However, most of them share a similar feature: it has not been resolved how these factors stimulate exocytosis. Moreover, a number of concerns about the bioenergetics of the formation of metabolic coupling factors have also been raised (Nicholls, 2016).

In sum, we have established an important role for [Ca^2+^]_mito_ in β-cell stimulus-secretion coupling. A novel mechanism, involving sequestering of Ca^2+^ into mitochondria, located at the vicinity of the voltage-dependent Ca^2+^-channels in the plasma membrane, that coupled with the well-established bioenergetic effects was unveiled (Figure 6); it accounts for the regulatory effect of mitochondrial Ca^2+^ uptake on insulin secretion. The recent molecular characterization of the mitochondrial calcium uniporter holocomplex holds promise for further elucidation of the complexities of β-cell stimulus-secretion coupling, in which the mitochondria are at centre stage.

**Figure 6.**
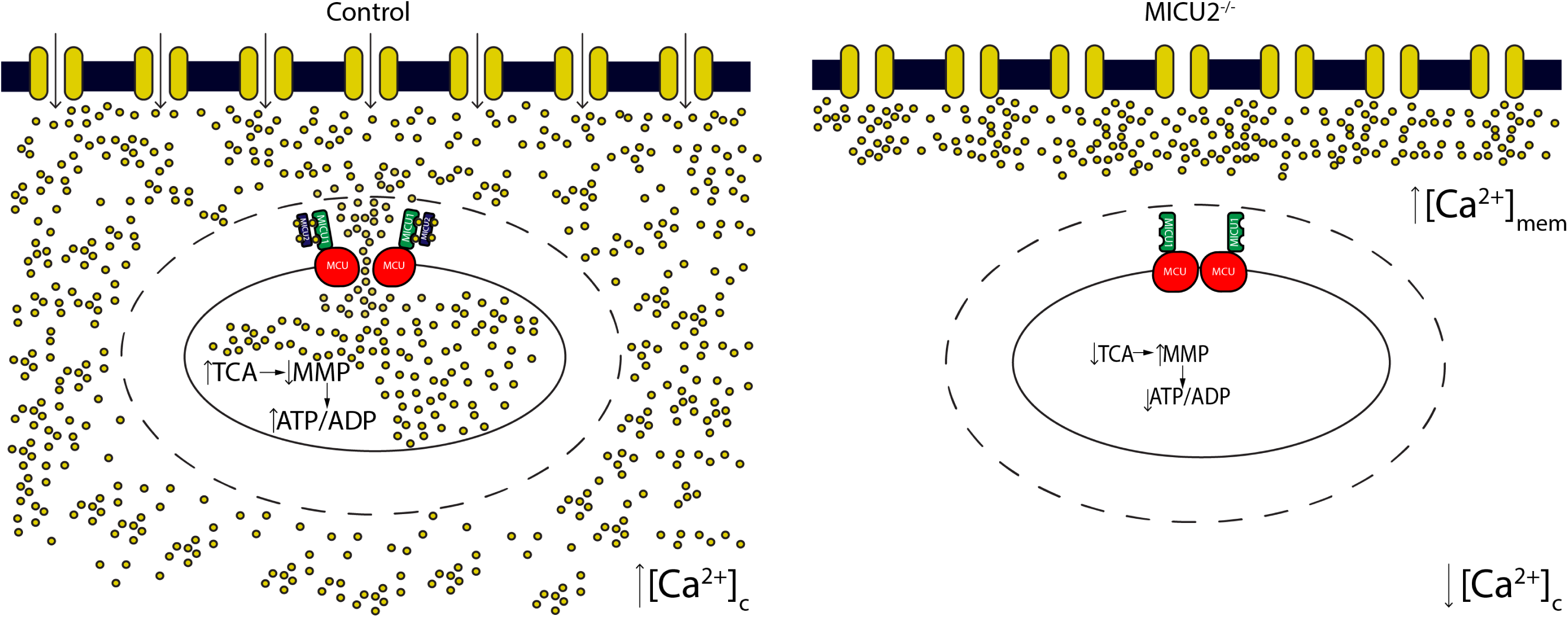
Model depicting role of mitochondrial Ca^2+^ uptake in cellular Ca^2+^ entry. Under control conditions, when mitochondrial Ca^2+^ uptake is normal, sub-plasmalemmal Ca^2+^ is cleared by cortical mitochondria. This prevents desensitization of voltage-gated Ca^2+^ channels, thereby facilitating an increase in cytosolic Ca^2+^. When *MICU2* is silenced, mitochondrial Ca^2+^ uptake is impaired. This prevents clearance of the sub-plasmalemmal Ca^2+^ layer, which inactivates voltage-gated Ca^2+^ channels, thereby reducing cytosolic Ca^2+^ elevation and consequently insulin secretion.

## EXPERIMENTAL PROCEDURES

### Cell Culture and Model Systems

INS-1 832/13 cells were cultured as previously described (Hohmeier et al., 2000). EndoC-βH1 cells (Ravassard, Hazhouz et al., 2011) were grown on Matrigel-fibronectin coated (100 μg/mL and 2 μg/mL, respectively, Sigma-Aldrich) culture vessels in DMEM containing 5.6 mM glucose, 2% BSA fraction V (Roche Diagnostics), 10 mM nicotinamide (Calbiochem), 50 μM 2-mercaptoethanol, 5.5 μg/mL transferrin, 6.7 ng/mL sodium selenite (Sigma-Aldrich), 100 U/mL penicillin, and 100 μg/mL streptomycin (PAA). Both cell lines were cultured at 37°C with 5% CO_2_. Unless otherwise stated, EndoC-βH1 cells were seeded at 2.3×10^5^ cells/cm^2^ and INS-1 832/13 cells at 1.5×10^5^ cells/cm^2^ in 24-well plates (Matrigel-fibronectin coated or uncoated) and transfected either with siRNA specific to *MICU2* or a scrambled siRNA. 48 hours after knock down, the growth media were changed to an overnight starvation medium containing 5.6 mM glucose for INS-1 832/13 cells and 1 mM glucose for EndoC-βH1 cells. HEK-293T cells were cultured in DMEM containing 1 g/L D-glucose, and supplemented with 10 % heat inactivated FBS and 100 U/mL penicillin, and 100 μg/mL streptomycin (PAA) (Pear, Nolan et al., 1993). Cells were plated at 1.5×10^5^ cells/cm^2^ in 24-well plates and transfected with either siRNA specific to *MICU2* or a scrambled siRNA. All assays were performed 48-72 hour after knock down.

### *Micu2* Knock Out Mice

We purchased a gene trap of *Micu2* from the Texas A&M Institute of Genomic Medicine (College Station, TX, USA), bred mice, and validated disruption of the *Micu2* gene, using standard procedures. The *Micu2* knock out mice are on a B57BL/6J background and have been extensively backcrossed. This mouse has been characterized previously (Bick et al., 2017).

### Insulin Secretion Assay

INS-1 832/13 cells were incubated in HEPES-balanced salt solution (HBSS; 115 mM NaCl, 5 mM KCl, 1 mM MgSO_4_, 1 mM CaCl_2_, 25.5 mM NaHCO_3_ 20 mM HEPES, 0.2% BSA, pH 7.2) containing 2.8 mM glucose for 2 hours followed by 1 hour of stimulation with HBSS containing 2.8 or 16.7 mM glucose (Hohmeier et al., 2000). Insulin secretion was measured with a rat insulin ELISA kit (Mercodia A/B, Sweden) according to manufacturer’s instructions. EndoC-βH1 cells were starved overnight in 1 mM glucose-containing growth medium a day before the experiment. On the day of the experiment, growth medium was removed and replaced with 1 mM glucose-containing HBSS for 2 hours followed by 1 hour of stimulation with HBSS containing 1 or 20 mM glucose (Scharfmann, Pechberty et al., 2014). Insulin secretion was measured with a human insulin ELISA (Mercodia A/B, Sweden) according to manufacturer’s instructions.

### RNA Isolation and Quantitative Real Time PCR

Total RNA was extracted from EndoC-βH1 cells, INS-1 832/13, and HEK-293T cells, using RNeasy Mini Kit (Qiagen) according to manufacturer’s protocol. RNA concentrations were determined using a NanoDrop Spectrophotometer (Thermo Scientific). A thousand ng total RNA were reverse transcribed using RevertAid First-Strand cDNA synthesis kit (Fermentas, Vilnius, Lithuania). Quantitative real time-PCR (qPCR) was performed using the TaqMan gene expression assays (*EFHA1/Micu2 rat*: s235193; *EFHA1/Micu2 human*: s47976; *Hprt1 rat*: Rn01527840; *Hprt1 human:* Hs99999909, Applied Biosystems, Life Technologies, Carlsbad, CA), using a 7900HT Fast Real-Time System (Applied Biosystems). The qPCR was carried out as follows. Five ng cDNA was mixed with 0.5 µL gene assay, 5 µL Taqman Mastermix and ddH_2_O to a final volume of 10 µL and qPCR was performed using the following program: 50 °C for 2 min, 95 °C for 10 min, 40 cycles of 95 °C for 15 s and 60 °C for 1 min. In all runs, a 10x dilution series was made of one of the samples and used to generate a calibration curve as well as quality control for amplification efficiency. Gene expression was relatively quantified using the calibration curve. The amount of mRNA was calculated relative to the amount of hypoxanthine-guanine phosphoribosyl transferase (*HPRT1*) as reference gene.

### SDS-PAGE and Western Blotting

Cells were harvested in RIPA buffer followed by lysis by shaking the samples for 30 minutes at 950 rpm and 4°C for 30 minutes. This was followed by centrifugation at 16000 g and 4°C for 20 minutes to remove cell debris. Supernatants/lysates were kept at −80⁰C until use. 10-20 μg protein were separated on mini-PROTEAN® TGX™ pre-stain gels (BIO-RAD) followed by activation of the pre-stain in BIO-RAD MP imager. Proteins were blotted onto Trans-Blot® Turbo^TM^ Transfer Pack (BIO-RAD) using BIORAD semidry blotter and total protein for loading control was determined from the pre-stain. Membranes were blocked with 10% BSA in TRIS-buffered saline (TBS) unless otherwise stated and hybridized with antibodies for MICU2/EFHA1 (Abcam, ab101465), β-tubulin (Abcam, ab6046) and α-Tubulin (Abcam, ab7291).

### Single Live Cell Cytosolic ATP and ATP/ADP Ratio Measurements Using Perceval and Perceval HR

Single cell cytosolic ATP/ADP ratio measurements were carried out in INS-1 832/13 cells using the genetically encoded biosensor Perceval HR (Tantama et al., 2013). Cells were seeded on poly-D-lysine coated 8-well chambered cover glasses (Lab-Tek, Thermo Scientific) at a density of 70,000 cells/cm^2^. Twenty-four hours after seeding, the cells were co-transfected with siRNA and 1 µg of plasmid encoding Perceval HR (Addgene ID:49083) at around 50% cell confluency. Cells were grown further for 48-72 hours before measurements. On the day of imaging, cells were pre-incubated at 37 °C in 400 μL buffer P (135 mM NaCl, 3.6 mM KCl, 1.5 mM CaCl_2_, 0.5 mM MgSO_4_, 0.5 mM Na_2_HPO_4_, 10 mM HEPES, 5 mM NaHCO_3_, pH 7.4) containing 2.8 mM glucose. After 1.5 hours of incubation, cells were imaged with 488 nm excitation and 505-535 nm emission recorded on a Zeiss LSM510 inverted confocal fluorescence microscope. Mouse islets were transduced with adenovirus expressing the similar ATP biosensor Perceval (Berg et al., 2009, Li, Shuai et al., 2013). Poly-D-lysine coated 8-well chambered cover glasses (Lab-Tek, Thermo Scientific) containing mouse islets were incubated for 2 hours with RPMI with 10% FBS and 100 U Penicillin and 100 mg/ml Streptomycin, containing the Perceval adenovirus. Then this medium was replaced with fresh RPMI medium with 10% FBS and 100 U Penicillin and 100 mg/ml Streptomycin and incubated overnight. On the day of the imaging, cells were pre-incubated at 37 °C in 400 μL buffer P (135 mM NaCl, 3.6 mM KCl, 1.5 mM CaCl_2_, 0.5 mM MgSO_4_, 0.5 mM Na_2_HPO_4_, 10 mM HEPES, 5 mM NaHCO_3_, pH 7.4) containing 2.8 mM glucose. After 1.5 hours of incubation, cells were imaged as described for the clonal cell lines with 488 nm excitation and emission recorded at 505-530 nm on a Zeiss LSM510 inverted confocal fluorescence microscope using a 40x/0.75 objective. Images were recorded at a frequency of 0.25 Hz (scan time = 3.93 s). For the 36 mM KCl solution, equimolarity was accounted for.

### Single Live Cell Cytoplasmic Free Ca^2+^ Measurements

Cells were seeded onto poly-D-lysine coated 8-well chambered cover glasses (Lab-Tek, Naperville, IL) at a density of 70,000 cells/cm^2^ per well. Twenty-four hours after seeding, the cells were transfected with siRNA and incubated for further 48-72 hours prior to imaging. On the day of imaging, cells were pre-incubated at 37 °C in 400 μL buffer P containing 2.8 mM glucose. After 1.5 hours incubation, based on variable Ca^2+^ affinities INS-1 832/13, EndoC-βH1 and HEK-293T cells were loaded with 2 μM of Fluo4 AM, Fluo8 AM, Fluo5F AM respectively. Following which incubation was continued for an additional 30 min. Immediately prior to imaging, the pre-incubation buffer was exchanged with 400 μL buffer P. Fluo4 AM, Fluo8 AM and Fluo5F AM were excited at 488 nm and emission recorded using a 505 nm emission filter on a Zeiss LSM510 inverted confocal fluorescence microscope using a 40x/0.75 objective. Images were recorded at a frequency of 0.64 hz (scan time = 1.57 s), using a pinhole diameter of 463 μm. Single cell free cytoplasmic Ca^2+^ traces were displayed in arbitrary fluorescent units.

For experiments performed with the linescan configuration, INS1 832/13 cells were plated, transfected and loaded as above. Imaging was again performed, using the Zeiss LSM510 inverted confocal fluorescence microscope using a 100x/1.45 objective with Fluo4 excited at 488 nm and emission recorded using a 505 nm emission filter. For each recording, a single line was drawn through the centre of 1-3 cells. This was the only part of the field of view that was measured during the recording. The pinhole diameter was reduced to 256 μm to minimise signal from outside of the focal plane (1.4 μm section). This allowed the Ca^2+^ signal to be tracked as it propagated from the edge to the centre of the cell with minimal interference from Ca^2+^ diffusing from above or below the focal plane. Recordings were performed at a frequency of 333 Hz (scan time = 3 ms) and a duration of 6000 cycles.

### Single Live Cell Mitochondrial Ca^2+^ Measurements

Single cell [Ca^2+^]_mito_ measurements were carried out using genetically encoded mitochondrial targeted Ca^2+^ biosensor mito-case12 (everogen cat# FP992.). Cells (INS-1 832/13 and HEK-293T) were seeded onto poly-D-lysine coated 8-well chambered cover glasses (Lab-Tek, Thermo Scientific) at a cell density of 70,000 cells/cm^2^. Twenty-four hours after seeding, the cells were co-transfected with siRNA and 1 µg of plasmid encoding mito-case12 at around 50 % cell confluency. The cells were further grown for 48-72 hours before measurements. On the day of imaging, cells were pre-incubated at 37 °C in 400 μL buffer P containing 2.8 mM glucose. After 1.5 hours cells were imaged using a 488 nm laser and emission recorded at 505-530 nm on a Zeiss LSM510 inverted confocal fluorescence microscope using a 100x/1.45 objective. Images were recorded at a frequency of 0.25 Hz (scan time = 3.93 s).

### Single Live Cell Mitochondrial Free Ca^2+^ Measurements using Rhod2 AM

EndoC-βH1 Cells were seeded onto poly-D-lysine coated 8-well chambered cover glasses at a density of 70,000 cells/cm^2^ (Lab-Tek, Naperville, IL). The day after seeding, cells were transfected with siRNA and incubated for 72 hours prior to imaging. On the day of imaging, cells were pre-incubated at 37 °C in 400 μL buffer P containing 2.8 mM glucose. After 1.5 hours incubation, 2 µM Rhod-2 AM (Abcam) were loaded for 30 min at 37 °C in buffer P. After this, cells were washed once with buffer P to remove excess dye prior to imaging. Excitation of Rhod-2 AM was performed at 543 nm and emission recorded at 590-615 nm on a Zeiss LSM510 inverted confocal fluorescence microscope. For single mouse pancreatic islet mitochondrial free Ca^2+^ measurement, poly-D-lysine coated 8-well chambered cover glasses containing single complete or partially digested mouse pancreatic islets pre-incubated at 37C in 400 μL buffer P containing 2.8 mM glucose. After 1.5 hours incubation, 2 µM Rhod-2 AM (Abcam) were loaded for 30 min at 37 °C in buffer P. After this, cells were washed once with buffer P to remove excess dye prior to imaging. Excitation of Rhod-2 AM was performed at 543 nm and emission recorded at 565-615 nm on a Zeiss LSM510 inverted confocal fluorescence microscope using a 100x/1.45 objective. Images were recorded at a frequency of 0.13 Hz (scan time = 7.86 s).

### Single Live Cell Mitochondrial Membrane Potential (Δψm) Measurements

INS-1 832/13 cells were seeded onto poly-D-lysine coated 8-well chambered cover glasses (Lab-Tek, Naperville, IL) at a density of 70,000 cells/cm^2^. Twenty-four hours after seeding, cells were transfected with siRNA and incubated for 48-72 hours prior to imaging. For Δψ_m_ measurements, cells were pre-incubated for 2 hours in buffer P with 100 nM TMRM (Invitrogen) and 2.5 µM cylosporin A. After incubation, cells were washed once with buffer P and then imaging was performed. Under these conditions, Δψ_m_ measurements were performed in quench mode (Nicholls & Ward, 2000). TMRM was present in the medium throughout the experiment. TMRM fluorescence measurements were performed using 543 nm excitation and a 560 nm long pass emission filter on a Zeiss LSM510 inverted confocal fluorescence microscope using a 40x/0..75 objective. Images were recorded at a frequency of 0.64 hz (scan time = 1.57 s).

### Single Live Cell Sub-plasmalemmal Ca^2+^ Measurements

Single cell [Ca^2+^]_mem_ measurements were carried out using pericam-based genetically encoded inner plasma membrane targeted Ca^2+^ probe mem-case12 (everogen cat# FP992.). Specifically, cells (INS1-832/13 and HEK 293T) were seeded onto poly-D-lysine coated 8-well chambered cover glasses (Lab-Tek, Naperville, IL) at a cell density of 70,000 cells/cm^2^. After 24 hours of seeding, the cells were co-transfected with siRNA and 1 µg of plasmid encoding mito-case12 at around 50% cell confluency. The cells were then grown further for 48-72 hours before measurements. On the day of imaging, cells were pre-incubated at 37 °C in 400 μL buffer P containing 2.8 mM glucose. After 1.5 hours cells were imaged with 488 nm excitation (8%) and emission recorded at 505-535 nm on a Zeiss LSM510 inverted confocal fluorescence microscope using a 100x/1.45 objective. Images were recorded at a frequency of 0.64 hz (scan time = 1.57 s).

### Mouse Islet Insulin Secretion

Islets were picked by hand as described earlier (Fex, Nitert et al., 2007) under the stereomicroscope and incubated in HBSS containing 2.8 mM glucose for 2 hours. This was followed by 1 hour of stimulation with HBSS supplemented with 2.8 or 16.7 mM glucose. Aliquots of incubation medium were collected and insulin secretion was measured with a mouse insulin ELISA kit (Mercodia A/B, Sweden) according to manufacturer’s instructions.

### Glucose and Insulin Tolerance Tests

All tolerance tests were performed in anaesthetized mice, using 0.25 mg midazolam (Midazolam Panpharma ®, Fougères, France) combined with 0.5 mg fluanison and 0.02 mg fentanyl (Hypnorm ®, VetPharma Ltd, Leeds, UK). For IVGTT, mice were starved for 2 hours prior to tail vein injection with 1g/kg D-glucose. For ITT, unfasted mice were intraperitoneally injected with 0.75 mU/g human insulin (Novo Nordisk, Clayton, NC, USA). In each case, blood was collected via retro-orbital bleed at the indicated time points post injection. Upon completion, samples were centrifuged, and plasma collected and stored at −20°C. Thereafter, insulin and glucose concentrations were determined by mouse insulin ELISA kit (Mercodia, Uppsala, Sweden) and Infinity™ Glucose (Oxidase) Reagent (Thermo Fisher Scientific, Waltham, MA, USA) respectively. Glucose measurements for ITT were determined by Amplex™ Red Glucose/Glucose Oxidase Assay Kit (Thermo Fisher Scientific, Waltham, MA, USA).

### Statistical analyses

All experiments were repeated on 3 different days with the exception of ITT, which was performed twice. Data in all figures are presented as Mean ± SEM, and unless specifically stated, statistical significance was determined by unpaired two-tailed Student’s t-test. For imaging experiments, n in figures represent number of cells or regions of interest (ROIs). For islet insulin secretion and IVGT assays, n in figures represent number of animals. For all statistical comparisons, *p*-value of less than 0.05 was considered significant.

## AUTHOR CONTRIBUTION

NV, AH, DGN and HM designed the study. NV, AH, AB, AMB, HB, YS, KK, PS, EC and MF performed the experiments. NV, AH, AMB and MF analysed the data. VM provided the *Micu2* knock out mice. AT, VM, DGN and HM gave critical insights to the project. NV, AH, AT, VM, DGN and HM wrote the manuscript.

## ACKNOWLEDGEMENT

Laila Jacobsson is acknowledged for genotyping mice. This study was supported by grants from the Swedish Research Council (14196-12-5, 2017-00956), EFSD/MSD, the Novo Nordisk foundation, Swedish Diabetes Foundation, The Swedish Strategic Research Foundation, Crafoord, Hjelt, Lars Hierta’s Minne, Fredrik och Ingrid Thuring’s, O.E. and Edla Johansson’s Vetenskapliga, Åke Wibergs, Direktör Albert Påhlssons, and Magnus Bergvalls Foundations, and the Royal Physiographic Society. This study was also supported by equipment grants from the KAW Wallenberg Foundation (2009-0243), and funding from the European Union’s Horizon 2020 Research and Innovation programme under grant agreement No 667191.

## STUDY APPROVAL

All experimental protocols for animals were approved by the local animal ethical committee of Lund/Malmö and complied with the guidelines of the Federation of European Laboratory Animal Science Association (FELASA).

